# Activity of *Plasmodium vivax* promoter elements in *Plasmodium knowlesi*, and a centromere-containing plasmid that expresses NanoLuc throughout the parasite life cycle

**DOI:** 10.1101/2021.03.30.437722

**Authors:** Roberto R. Moraes Barros, Kittisak Thawnashom, Tyler J. Gibson, Jennifer S. Armistead, Ramoncito L. Caleon, Miho Kaneko, Whitney A. Kite, J. Patrick Mershon, Jacqueline K. Brockhurst, Theresa Engels, Lynn Lambert, Sachy Orr-Gonzalez, John H. Adams, Juliana M. Sá, Osamu Kaneko, Thomas E. Wellems

## Abstract

**Background:** *Plasmodium knowlesi* is now the major cause of human malaria in Malaysia, complicating malaria control efforts that must attend to the elimination of multiple *Plasmodium* species. Recent advances in the cultivation of *P. knowlesi* erythrocytic-stage parasites *in vitro*, transformation with exogenous DNA, and infection of mosquitoes with gametocytes from culture have opened up studies of this pathogen without the need for resource-intensive and costly non-human primate (NHP) models. For further understanding and development of methods for parasite transformation in malaria research, this study examined the activity of various trans-species transcriptional control sequences and the influence of *Plasmodium vivax* centromeric (*pvcen*) repeats in plasmid-transfected *P. knowlesi* parasites.

**Methods:** *In vitro* cultivated *P. knowlesi* parasites were transfected with plasmid constructs that incorporated *P. vivax* or *Plasmodium falciparum* 5’ UTRs driving the expression of bioluminescence markers (firefly luciferase or Nanoluc). Promoter activities were assessed by bioluminescence, and parasites transformed with human resistant allele dihydrofolate reductase-expressing plasmids were selected using antifolates. The stability of transformants carrying *pvcen*-stabilized episomes was assessed by bioluminescence over a complete parasite life cycle through a rhesus macaque monkey, mosquitoes, and a second rhesus monkey.

**Results:** Luciferase expression assessments show that certain *P. vivax* promoter regions, not functional in the more evolutionarily-distant *P. falciparum*, can drive transgene expression in *P. knowlesi*. Further, *pvcen* repeats may improve the stability of episomal plasmids in *P. knowlesi* and support detection of NanoLuc-expressing elements over the full parasite life cycle from rhesus macaque monkeys to *Anopheles dirus* mosquitoes and back again to monkeys. In assays of drug responses to chloroquine, G418 and WR9910, antimalarial half-inhibitory concentration (IC_50_) values of blood stages measured by NanoLuc activity proved comparable to IC_50_ values measured by the standard SYBR Green method.

**Conclusion:** All three *P. vivax* promoters tested in this study functioned in *P. knowlesi* whereas two of the three were inactive in *P. falciparum*. NanoLuc-expressing, centromere-stabilized plasmids may support high-throughput screenings of *P. knowlesi* for new antimalarial agents, including compounds that can block the development of mosquito- and/or liver-stage parasites.

## Background

*Plasmodium knowlesi*, once thought to infect macaques almost exclusively, is now a major cause of human malaria in Malaysia (1). Up to 10 % of *P. knowlesi* human infections cause severe malaria, leading to death in 1-2% of those afflicted (2). Increased prevalence of *P. knowlesi* infections represents an additional challenge to malaria elimination, especially in Southeast Asia, where *Plasmodium falciparum* and *Plasmodium vivax* infections are in decline (3,4). In some regions endemic to multiple *Plasmodium* species, malaria control work was found to decrease the prevalence of *P. falciparum* but did not produce comparable decreases of *P. vivax* or *P. knowlesi* (3-5).

In the 1970s, laboratory research on *P. falciparum* was boosted by the development of *in vitro* cultivation methods (6,7). However, similar long-term cultivation of any other human malaria species was unavailable until the adaptation of *P. knowlesi* parasites to culture (8,9). As with *P. falciparum*, transfection methods and the ability to infect mosquitoes with cultivated *P. knowlesi* parasites followed (8–11), and these are supporting new experimental studies without the need for resource-intensive and costly non-human primate (NHP) models. Fundamental biological differences between the *Plasmodium* species and factors affecting malaria control that are now more accessible to investigation include: the period of the intraerythrocytic life cycle (24 h for *P. knowlesi vs*. 48 h for *P. falciparum*) (12,13); features of host cell preference and cell invasion mechanisms (14–16); gametocyte development and transmission to mosquitoes (17); and antimalarial drug responses and mechanisms of resistance (18,19).

Transfection and genetic transformation of malaria parasites with various bioluminescent proteins have been used for *in vivo* studies of the traversal and tissue invasion of sporozoites injected in mammalian hosts by mosquitoes (20); functional analyses of promoters and introns (21–25); screens for candidate anti-malarials and inhibitors of parasite growth (26–28); and evaluations of compounds that block parasite infectivity to mosquitoes (29). This report presents investigations of exogenous transcriptional control sequences in transfected *P. knowlesi* parasites. Results show that *P. knowlesi* parasites can recognize *P. vivax* promoters that are poorly recognized by *P. falciparum* parasites; further, *P. vivax* centromeric (*pvcen*) repeats can improve the stability of transgenic plasmids across the complete parasite life cycle of *P. knowlesi* in rhesus monkeys and *Anopheles dirus* mosquitoes.

## Methods

### Parasites and culture

*P. falciparum* parasites were maintained in human red blood cells (hRBCs) purchased from Interstate Blood Bank (Memphis, TN, USA). *P. knowlesi* parasites were cultivated in rhesus red blood cells (rRBCs) as described (10); blood samples were obtained according to the National Institutes of Health (NIH) Guidelines for Animal Care and Use, under an Animal Study Proposal (ASP) approved by the National Institute of Allergy and Infectious Diseases (NIAID) Animal Care and Use Committee (ACUC); rhesus blood was processed and plasma was removed as described (10).

Cryopreserved stocks of a *P. falciparum* 3D7 clone (30) and the *P. knowlesi* H strain (31) were obtained from NIH inventory and thawed by standard methods (32). *Plasmodium falciparum* cultures were propagated in *P. falciparum* complete RPMI (PF-cRPMI) consisting of RPMI-1640 (KD Medical, Columbia, MD, USA) supplemented with 25 mM HEPES, 50 μg/mL hypoxanthine, 0.21% sodium bicarbonate, 20 mg/L gentamicin, and 1% Albumax II (Life Technologies, Carlsbad, CA, USA). *Plasmodium knowlesi* cultures were maintained in *P. knowlesi* complete RPMI (PK-cRPMI), consisting of RPMI-1640 supplemented with 25 mM HEPES, 50 μg/mL hypoxanthine, 0.26% sodium bicarbonate, 10 mg/L gentamicin (KD Medical), and 1% Albumax II (32). Cultures were maintained with RBCs up to 5% hematocrit, under a 90% N_2_/5% CO_2_/5% O_2_ gas mixture at 37°C with daily media changes. Parasite development and propagation across the asexual blood-stage cycle were monitored by microscopy of thin blood films after fixation in methanol and staining for 15 min with 20% Giemsa’s solution (Sigma-Aldrich, St. Louis, MO, USA). Parasitemia was estimated by counting the number of parasitized RBCs (pRBCs) per 1,000 RBCs. Cultures were maintained at parasitemias between 0.5 and 10%.

### Rhesus infection using blood-stage *P. knowlesi*

Rhesus macaques (*Macacca mulatta*) were obtained from NIH-approved sources and housed in compliance with the Animal Welfare Act and the Guide for the Care and Use of Laboratory Animals (ILAR, 1996). All animal care and use in this study were performed in accordance with the NIH Animal Research Advisory Committee (NIH ARAC) Guidelines, under ASP protocols approved by the NIAID ACUC.

Animals were anesthetized intramuscularly (IM) with 10 mg/kg ketamine to allow infection by mosquito bites or by intravenous (IV) inoculation of cultivated blood stage parasites. For the IV inoculation, a culture sample containing ~2 × 10^8^ parasitized rRBCs was pelleted by centrifugation (700 x g, 3 minutes). Pelleted cells were washed with 10 mL sterile and pure RPMI-1640, and resuspended to 1.5 mL volume of the same medium. After infection, parasitemia was checked daily by Giemsa-stained thin blood films as described above.

*Plasmodium knowlesi* infections were cured with three daily administrations of oral chloroquine (total dose: 50 mg/kg body weight). Thin and thick blood films were checked up to 30 days after treatment to confirm cure.

### Mosquito feedings and assessment of oocysts and sporozoites

Mosquito feedings were performed using 3 – 5-day-old female *Anopheles. dirus* B and *An. dirus* (X strain) mosquitoes (31) raised in the LMVR/NIAID insectary. Mosquitoes were starved for 24 h in secure pint containers with up to 100 female mosquitoes per container. Feedings were performed at night (12:00 AM), as recommended for *P. knowlesi* transmission (12). Two pints of mosquitoes were used in each experiment, and mosquitoes were allowed to feed through a double safety net (bridal veil) for ~20 min directly onto the inner arm or thighs of the anesthetized rhesus macaque. Fed mosquitoes were maintained with 10% corn syrup in water at 26°C for up to 18 days in the LMVR human malaria-secure insectary.

Mosquitoes were randomly selected and dissected 6 – 9 days post-feeding and examined for oocysts in the midgut; additional mosquitoes were randomly selected and dissected after 10 days to examine for sporozoites in the salivary glands. To assess oocyst presence and number, midguts from 16 – 20 dissected mosquitoes per container were stained with 0.05% mercurochrome for 10 min and then examined under the microscope (20× – 40× objective lens). Individual mosquito midguts containing oocysts were transferred to 1.5 mL tubes and used for NanoLuc assays. Sporozoites isolated from dissected salivary glands were combined and counted using a hemocytometer under the microscope at 40× magnification. The sporozoites were transferred to 1.5 mL tubes and used for NanoLuc assays. Remaining infected mosquitoes were used 14 days post-feeding to transmit parasites to an uninfected rhesus macaque as described above.

### Plasmids

Plasmid pD-pfcam-Luc (10) contains the *fLuc* (firefly luciferase) cassette under control of a 0.6 kb 5’ untranslated region (UTR) from *P. falciparum calmodulin* (*pfcam*) and a 0.8 kb 3’ UTR sequence from *P. falciparum heat shock protein 86 (pfhsp86)*. pD-pfcam-Luc also contains the *hdhfr* cassette under control of a 0.6 kb 5’ UTR from *Plasmodium chabaudi dhfr-ts* (*pcdts* 5’), and a 0.8 kb 3’ UTR from *P. falciparum histidine rich protein 2* (*pfhrp2*) **(Fig 1)**.

**Figure 1.**
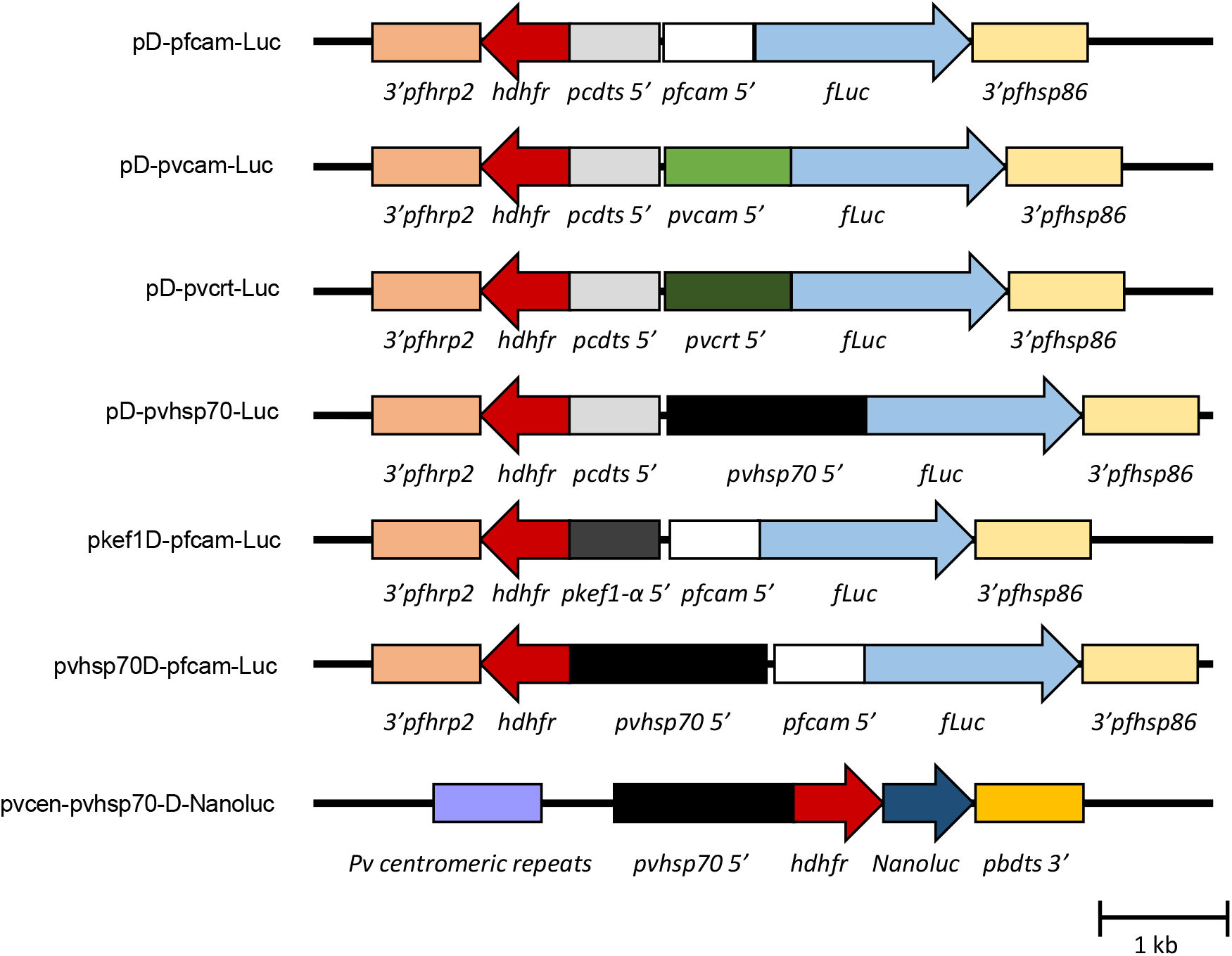
Plasmid constructs used to evaluate *Plasmodium* promoter sequences in *P. knowlesi*. Expression of *hdhfr* from the plasmids confers resistance to WR99210 and a *fLuc or NanoLuc* reporter. The pD-pfcam-Luc plasmid has the *fLuc* cassette driven by the *P. falciparum calmodulin* 5’ UTR (*pfcam* 5’) and the *hdhfr* cassette driven by the *P. chabaudi dts* 5’ UTR (*pcdts* 5’). pD-pvcam-Luc, pD-pvcrt-Luc and pD-pvhsp70-Luc have the 5’ UTR sequences of *P. vivax calmodulin* (*pvcam* 5’), *chloroquine resistance transporter* (*pvcrt* 5’) and *heat shock protein 70* (*pvhsp70* 5’) driving *fLuc* expression, respectively, replacing the *pfcam* promoter from the pD-pfcam-Luc. Plasmids pvhsp70D-pfcam-Luc and pkef1D-pfcam-Luc have the 5’ UTR sequences of *P. vivax heat shock protein 70* (*pvhsp70* 5’) and *P. knowlesi elongation factor 1 a* (*pkef1-α* 5’), respectively, replacing the *pcdts* promoter from the pD-pfcam-Luc to drive the expression of *hdhfr*. The pvcen-pvhsp70-D-NanoLuc plasmid includes *P. vivax* centromeric sequence repeats from chromosome 11 (*Pv centromeric repeats*) and has *hdhfr-NanoLuc* fusion expression driven by the *P. vivax heat shock protein 70* 5’ UTR (*pvhsp70* 5’). The arrows indicate directions of transcription.

*P. vivax* and *P. knowlesi* 5’ UTR sequences were amplified from genomic DNA of *P. vivax* NIH-1993 (33) or *P. knowlesi* H strains using the primers listed in Supplementary Table 1. Amplified sequences of 0.7 kb from the *P. vivax calmodulin* (*pvcam*) 5’ UTR, 0.9 kb of the *P. vivax chloroquine resistance transporter* (*pvcrt*) 5’ UTR, 1.4 kb of the *P. vivax heat shock protein 70* (*pvhsp70*) 5’ UTR, and 0.6 kb of the *P. knowlesi elongation factor-1 alpha* (*pkef1-alpha)* 5’ UTR were isolated from agarose gels and cloned into the pGEM-T vector (Promega, Madison, WI, USA). To replace the *pfcam* 5’ UTR sequence that drives *luciferase* transcription in the pD-pfcam-Luc, the pGEM-T clones of the *pvcam, pvcrt*, and *pvhsp70* 5’ UTRs were digested using BlpI and SpeI restriction enzymes (New England Biolabs, Ipswich, MA), and agarose gel-purified fragments were subcloned into the plasmid pD-pfcam-Luc also digested with these enzymes. Resultant plasmids were named pD-pvcam-Luc, pD-pvcrt-Luc and pD-pvhsp70-Luc. To replace the *P. chabaudi dhfr-ts* 5’ UTR that drives the *hdhfr* transcription in the pD-PfCam-Luc, the pGEM-T clones of the *pvhsp70* and *pkef1-alpha* 5’ UTRs were digested using BlpI and NcoI, and the resulting fragments were cloned into the plasmid previously digested with the same enzymes, thus generating pvhsp70D-pfcam-Luc and pkef1D-pfcam-Luc.

Plasmid pvcen-pvhsp70-D-NanoLuc was developed from pDST-PvCEN11S3, pENT41-PvHSP70-5U-1k, pENT12-hDHFR, and pENT23-NLuc by MultiSite Gateway LR reaction (Invitrogen, Carlsbad, CA, USA) (34). pENT23-NLuc was generated by Gateway BP reaction between pDONR P2R-P3 plasmid (Invitrogen) and a DNA fragment encoding NanoLuc Luciferase amplified from pNL1.1 (NanoLuc) vector (Promega) with primers Nluc.B2F and Nluc.B3R **(Supplementary Table 1).** The plasmid contains the centromeric sequences of *P. vivax* chromosome 11 and harbors a 1 kb 5’ UTR sequence from the *P. vivax heat shock protein* driving the expression of a hdhfr-NanoLuc fusion cassette **(Fig 1)**. The plasmid also contains 800 bp of the 3’ UTR sequence from the *Plasmodium berghei* bifunctional dihydrofolate reductase-thymidylate synthase gene.

### Parasite transformation

Parasites were transformed by the spontaneous DNA uptake method as described (10,35). Briefly, 300 μL of pelleted rhesus RBCs were washed twice by centrifugation and resuspended in 5 mL cytomix (120 mM KCl, 0.15 mM CaCl, 2 mM EGTA, 5 mM MgCl_2_, 10 mM K_2_HPO_4_/KH_2_PO_4_, 25 mM Hepes, pH adjusted to 7.6 with KOH). The RBCs were recovered and mixed with 100 μL of cytomix containing 40 μg of plasmid DNA, and transferred to a 0.2 cm electroporation cuvette (Bio-Rad, Hercules, CA, USA). Uninfected RBCs were electroporated in a Bio-Rad Gene Pulser II at 310 V and 975 μF, with resulting time constants of 20-35 msec. The plasmid-loaded RBCs were washed twice with 5 mL of incomplete RPMI (cRPMI without Albumax II) and used immediately. Plasmid-loaded RBCs were added to 10 mL of PK-cRPMI in a 25 cm^2^ flask. A 50 μL suspension of 3% asynchronous pRBCs at 5% hematocrit was added to the flask for a final parasitemia of 0.5%. After 24 h in culture, fresh plasmid-loaded RBCs were again added to the flask for *P. knowlesi* transfections. After 72 h in culture, cells were collected for luciferase assays and/or continued in culture under 1 nM WR99210 pressure (WR99210 kindly supplied by Jacobus Pharmaceuticals Co., Inc.).

### Luciferase assays

The fLuc and NanoLuc enzymatic assays were performed using blood- and mosquito-stage parasites. To remove hemoglobin from RBCs and avoid a possible quenching effect of luciferase assay signal, RBCs were treated with saponin: up to 250 μL of packed RBCs were pelleted from cultures by centrifugation (800× *g*, 3 min), overlying supernatant culture medium was removed, and the cells were twice suspended and repelleted in 1,000 μL of 0.15% saponin in PBS (10 mM PO_4_^3-^, 137 mM NaCl, 2.7 mM KCl). After washing twice with 1 mL of PBS, the parasites were pelleted, resuspended in 50 μL (fLuc assays) or 100 μL (NanoLuc assays) of 1× Cell Culture Lysis Reagent (Luciferase Assay System; Promega) and incubated for 10 min at room temperature. The resulting lysate was centrifuged for 1 min at 10,600× *g* and the supernatant (20 μL upper liquid layer) was used for the enzymatic reaction.

Midguts containing *P. knowlesi* oocysts were isolated from *Anopheles* mosquitoes. Each isolated midgut was re-suspended in 20 μL of PBS, mixed with 80 μL of Passive Lysis Buffer (Promega) and incubated for 10 min at room temperature for cell lysis. The lysate was centrifuged for 1 min at 10,600× *g* and the supernatant (upper liquid layer) was used for the NanoLuc assays. Sporozoites isolated from 8 salivary gland pairs were combined and re-suspended in 40 μL of PBS. The sporozoites were mixed with 60 μL of Passive Lysis buffer and incubated for 10 min at room temperature for cell lysis. After centrifugation (1 min at 10,600× *g*) the supernatant was diluted 10-fold in Passive Lysis Buffer and the dilution was utilized for NanoLuc assays.

Firefly luciferase assays were performed using the Luciferase Assay System (Promega) following manufacturer’s instructions. For this assay, 20 μL of the lysate supernatant was mixed with 100 μL of Promega Luciferase Assay Reagent and luminescence was assessed with a GloMax 20/20 Luminometer (10 sec integration time). NanoLuc assays were performed using the Nano-Glo Luciferase Assay System (Promega) following manufacturer’s instructions. For the enzymatic reaction, 40 μL of the lysate supernatant were mixed with the same volume of Nano-Glo reaction solution (Promega), incubated for 3 min at room temperature, and assessed with a GloMax 20/20 Luminometer (1 sec integration time). Measurements were collected in technical duplicates and averaged for analysis.

### Quantitative Real-Time PCR

Quantitative real-time PCR (qPCR) reactions to evaluate plasmid copy number were performed using the Rotor-Gene SYBR Green PCR Kit according to manufacturer’s instructions (Qiagen, Hilden, Germany). For each reaction a total of 50 ng of genomic DNA and 200 nM of each oligonucleotide primer were mixed with the Rotor-Gene PCR SYBR Green mix and the reaction was performed in the Rotor-Gene Q instrument, following the PCR cycle: 10 sec at 96°C, 30 sec at 60°C, repeated 35 times. Primers NanoLuc RT-F-2 and NanoLucRT-R-2 were designed to amplify a 133 bp sequence of the NanoLuc sequence. Primers Pk-Aldolase-F-2 and Pk-Aldolase-R were designed to amplify a 147 bp sequence of the single copy gene *Pk aldolase* (access number: PKNH_1237500.1). The threshold cycle value (Ct) for each gene fragment was used to estimate the ΔCt of each sample compared with that of the single copy gene *Pk aldolase*, and the ΔΔCt (36) was used to calculate relative proportions of NanoLuc copies in the different samples.

The samples used in this analysis were collected from *in vitro* cultures containing 1 nM WR99210 (Jacobus Pharmaceuticals, Princeton, NJ) for selective pressure, without drug pressure for 14 days, and from two infected rhesus macaques (rhesus #1 and rhesus #2). One sample was collected at day four of the first infection (rhesus #1) and the second sample was collected at day six of the second infection (rhesus #2). The *in vitro* culture sample under drug pressure was used as reference sample.

### Antimalarial half-maximum inhibitory concentration (IC_50_) determinations

Blood-stage *P. knowlesi* cultures were diluted to 0.5% parasitemia and brought to 2% hematocrit for growth in 96-well plates containing the desired drug concentration series (200 μL of culture/well). Plates were incubated for 40 h at 37°C in a humidified chamber containing 5% CO_2_, 5% O_2_, and 90% N_2_. Parasite growth in the plates was determined by quantification of double stranded DNA in each well (SYBR Green method) (37) or by the luminescence measured in each well (NanoLuc method).

For the SYBR Green method, the plates were removed from the incubator and frozen overnight. DNA amounts representing parasite growth were quantified by the SYBR Green I intercalating fluorescent dye (Life Technologies) as described (38). Plates were protected from light during the dye incubation period (30 – 60 min) and fluorescence intensities were measured using a FLUOstar Optima Microplate Reader (BMG Labtech, Ortenberg, Germany) with excitation and emission settings of 485 nm and 535 nm, respectively.

For the NanoLuc method, plates were removed from the incubator and 100 μL of NanoGlo IC_50_ solution (Promega NanoGlo reaction mixture diluted 10-fold in PBS) were added to each well and mixed by pipetting. The plates were then incubated for 3 min at room temperature and the luminescence was assessed using a plate luminometer (Molecular Devices, San Jose, CA, USA) with 1 sec integration time. Fluorescence and luminescence readings were normalized to values from control wells containing no drug. IC_50_ values were determined from fitted response curves (non-linear regression with variable slope, GraphPad Prism Software), and data from at least two independent assays were used to calculate the average IC_50_ value of the *P. knowlesi* pvcen-pvhsp70-D-NanoLuc transgenic line with each method.

### *Plasmodium knowlesi* red blood cell invasion assays

*P. knowlesi* tightly synchronized schizonts were obtained by magnetic purification using MACS column (Miltenyi Biotec, Bergisch Gladbach, Germany) (39), adjusted to a 0.5% parasitemia in the target RBCs and incubated for 16 h at standard culture conditions. Rhesus RBCs were used as positive controls and mouse RBCs as negative controls. Invasion of human, *Aotus nancymaee*, and *Saimiri boliviensis* RBCs was evaluated in the experiments. Invasion was determined by counting the number of ring-infected RBCs per 5,000 RBCs (microscopy counts), and expressed relative to the counts observed in rhesus RBCs (positive control). Assay statistics were obtained from at least three independent experiments. For NanoLuc evaluation of RBC invasion, pvcen-pvhsp70-D-NanoLuc-transformed parasites were used for the invasion assays and the NanoLuc activity was assessed using the NanoGlo 20/20 luminometer as described above.

## Results

### *P. vivax* 5’ UTR are broadly active in *P. knowlesi* but limited in *P. falciparum*

To evaluate the cross-species activity of different *Plasmodium* gene regulatory sequences, a variety of plasmids were constructed for expression of firefly luciferase (*fLuc*) under control of *P. vivax* or *P. falciparum* 5’ UTR sequences. Plasmids pD-pvcam-Luc, pD-pvcrt-Luc and pD-pvhsp70-Luc contain 5’ UTR sequences from the genes for *P. vivax* calmodulin (*pvcam*), chloroquine resistance transporter (*pvcrt*), and heat shock protein 70 (*pvhsp70*), respectively **(Fig 1)**. Plasmid pD-pfcam-Luc **(Fig 1)** contains a 5’ UTR sequence of the *P. falciparum* calmodulin gene (*pfcam*) and is a flanking region previously found to have cross-species promoter activity in *P. knowlesi* (10).

For measurements of luciferase activity, pRBC were taken from asynchronous *P. falciparum* or *P. knowlesi* cultures and transformed by introduction to culture with plasmid-loaded RBCs (spontaneous DNA uptake method) (10,35). **Fig 2** shows the relative luciferase levels from the different transfected parasite cultures normalized to the levels from parasites transfected with the plasmid pD-pfcam-Luc. Each of the four plasmid constructs yielded signals in *P. knowlesi* ranging from 0.5 to 4.4 × of the pD-pfcam-Luc signal. In contrast, only two of the four plasmids provided luciferase signals in *P. falciparum*; no signals were obtained from *P. falciparum* parasites transfected with the *P. vivax* promoter constructs pD-pvcam-Luc and pD-pvcrt-Luc **(Fig 2, Supplementary Table 2)**. These results are consistent with a previous report suggesting that *P. falciparum* is unable to recognize certain *P. vivax* promoters for expression (22). Nevertheless, *P. falciparum* parasites transfected with the plasmid pD-pvhsp70-Luc presented luciferase levels similar to those from *P. knowlesi* transformants, 4.55 ± 0.18–fold higher (average ± standard error of the mean) than the level from parasites transfected with the control plasmid pD-pfcam-Luc.

**Figure 2.**
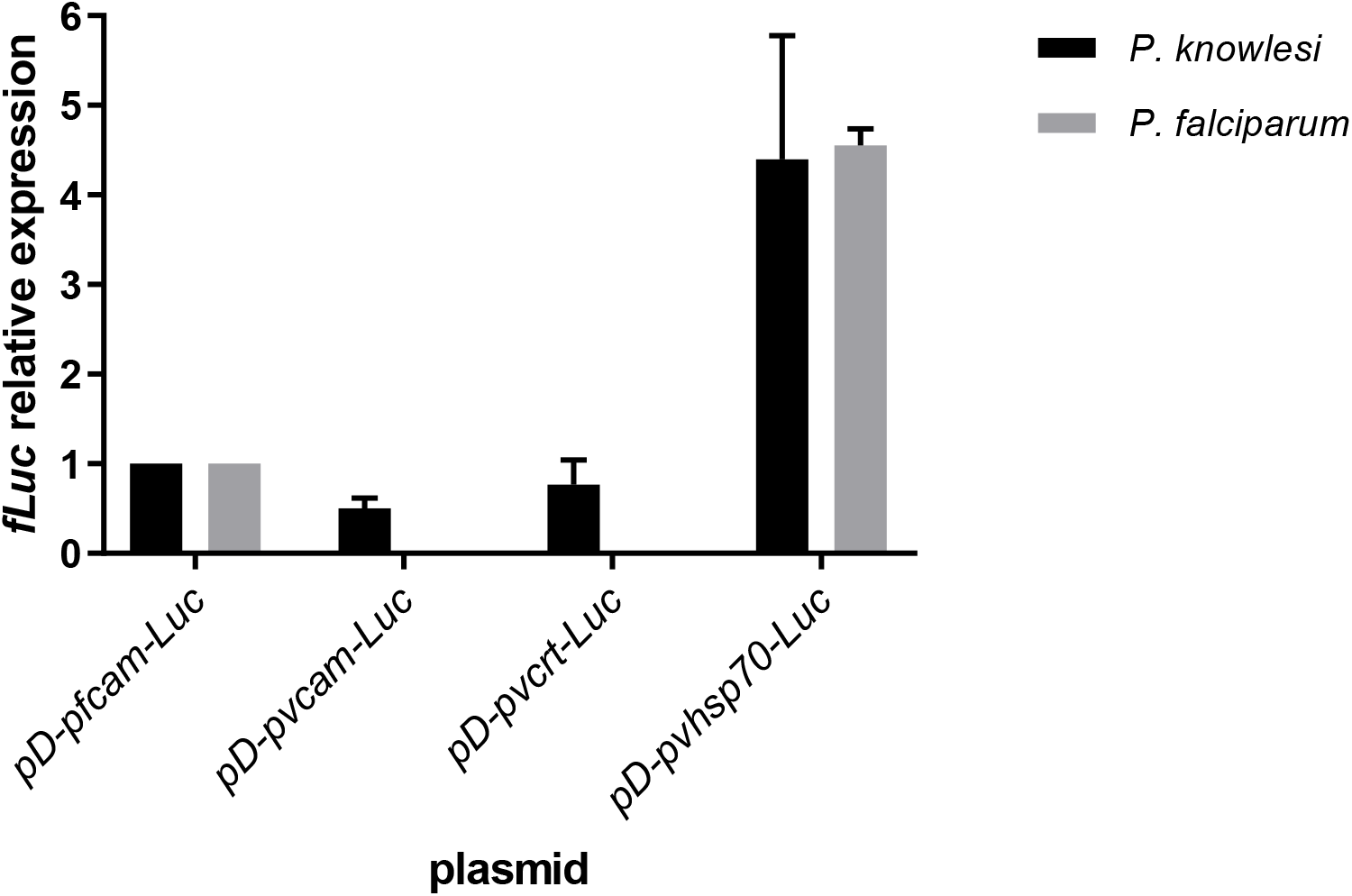
*P. knowlesi* recognizes *P. vivax* promoter regions not recognized by *P. falciparum*. *P. falciparum* and *P. knowlesi* parasites were transformed *in vitro* by spontaneous DNA uptake from RBCs pre-loaded with plasmids pD-pfcam-Luc, pD-pvcam-Luc, pD-pvcrt-Luc, or pD-pvhsp70-Luc. Luciferase activity measurements were obtained after 72 h of parasite cultivation in the plasmid-loaded RBCs and were normalized relative to the activity obtained from parasites transformed with pD-pfcam-Luc. Values represent the mean ± standard error from three independent experiments.

The *P. vivax hsp70* promoter sequence was also tested in *P. knowlesi* for expression of a drug resistance sequence used in selection of transgenic parasites. **Figure 1** presents schematics of the plasmids for this purpose: pvhsp70D-pfcam-Luc contains the *pvhsp70* promoter sequence driving *hdhfr* expression that confers resistance to antifolates such as WR99210 and pyrimethamine; pkef1D-pfcam-Luc contains the *pkef1-alpha* promoter previously used to drive *hdhfr* expression for selection of transgenic parasites *in vitro* (9). Both plasmids contain the *fLuc* reporter sequence under control of the *pfcam* promoter **(Fig 1)**. **Fig 3A** shows the parasite growth observed by thin blood film microscopy in the cultures transfected with the different plasmids. Parasites were not detected by thin blood films in either culture 4 days after the start of drug selection with 1 nM WR99210, but they became detectable on day 14 (parasitemias = 0.1%) **(Supplementary Table 3)**. **Fig 3B** shows the luciferase activity obtained from 80 μL of RBCs collected from the cultures under selection. The luminescence signal of pvhsp70D-pfcam-Luc transgenic parasites increased an average of 1.34–fold per day for the first 12 days under selection, similar to the pkef1D-pfcam-Luc culture (average increase of 1.46-fold/ day) **(Fig 3B, Supplementary Table 3)**. Over a period of three weeks of drug pressure, the transformed parasites were observed to grow at a rate similar to untransfected cultures *in vitro* (3 – 5-fold/day; data not shown).

**Figure 3.**
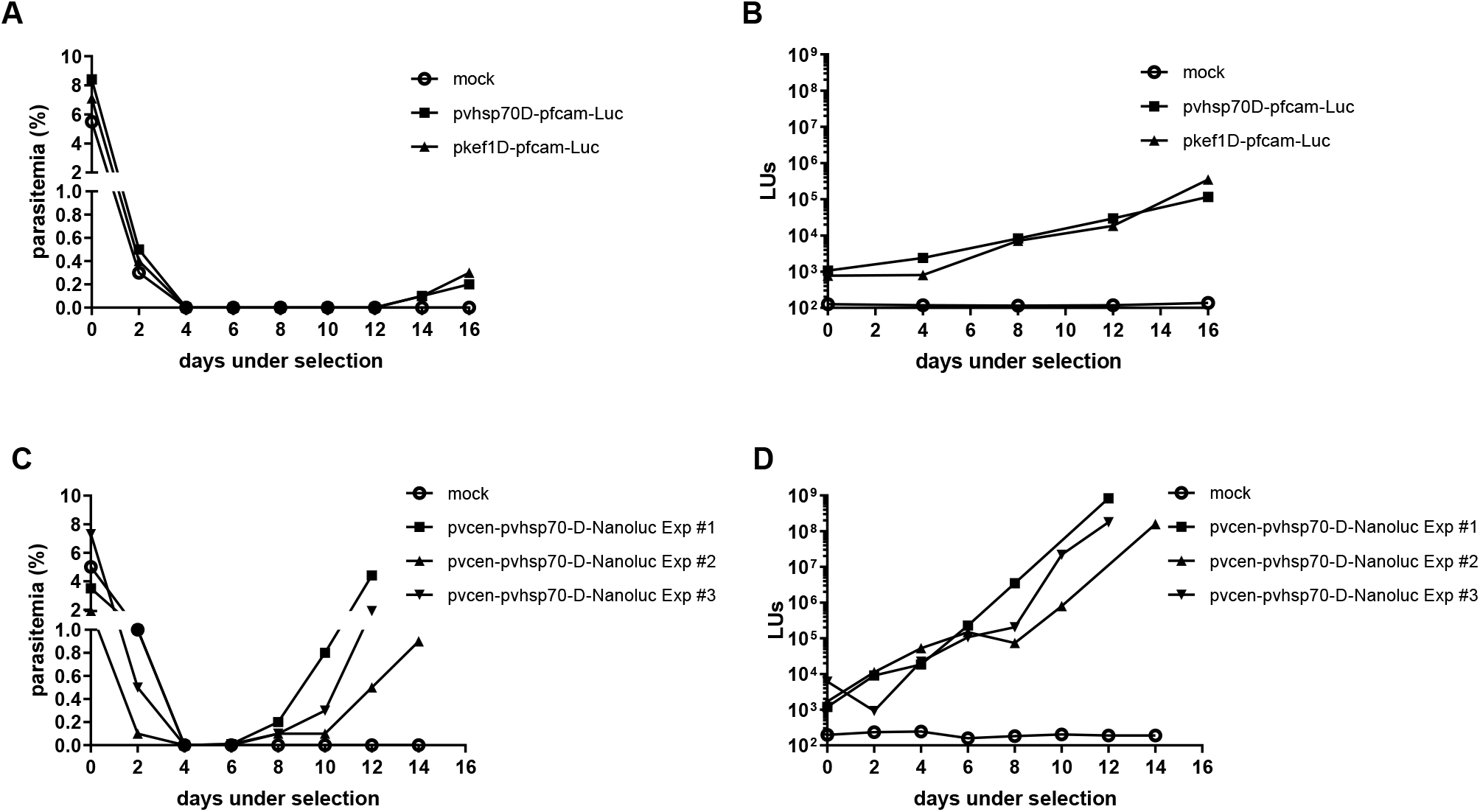
Selection of drug-resistant *P. knowlesi* harboring markers under the control of *P. vivax* regulatory sequences. Drug selection with 1 nM WR99210 was initiated 72 h after addition of plasmid-loaded RBCs to parasite cultures. (A) Parasitemia counts by microscopy of cultures transfected with plasmids pvhsp70D-pfcam-Luc and pkef1D-pfcam-Luc over 16 days of drug selection. (B) Luminescence measurements from samples of the transformant cultures presented in Fig 3A. (C) Parasitemia counts by microscopy of cultures transfected with plasmid pvcen-pvhsp70-D-NanoLuc in three independent experiments. (D) Luminescence measurements from samples of the transformant cultures presented in Fig 3C. LUs, luminescence units; mock, control transformation experiments performed in parallel without plasmid; parasitemia, percentage of RBCs infected with *P. knowlesi* parasites (counts of 1,000 RBCs).

### Rapid selection of transgenic parasites containing plasmids with *P. vivax* centromeric sequences

The inclusion of centromeric sequences in plasmids and artificial chromosomes can support maintenance of exogenous DNA in transfected parasites, with proper segregation through the multiple stages of their life cycle. Examples of success with this strategy have been reported with *P. berghei, P. falciparum*, and *Plasmodium cynomolgi* (40,41). To improve the maintenance and segregation of a selectable marker with very high sensitivity in detection assays, a plasmid containing *P. vivax* centromeric sequences (*pvcen*) was developed and tested in *Plasmodium yoelli* (34). Here, this plasmid was modified to generate pvcen-pvhsp70-D-NanoLuc, which expresses a fused-NanoLuc reporter cassette driven by the *P. vivax hsp70* promoter sequence **(Fig 1).**

*P. knowlesi* parasites were transfected with pvcen-pvhsp70-D-NanoLuc by the spontaneous DNA uptake method and drug-resistant parasites were selected by 1 nM WR99210 exposure. In three independent experiments, transgenic parasites containing the centromeric plasmid were detected after 8 days of drug pressure (parasitemia ≥ 0.1%) **(Fig 3C, Supplementary Table 4),** 6 days earlier than the detection of transgenic parasites containing episomal constructs without the centromeric sequences **(Fig 3A, Supplementary Table 3)**. In luminescence assays, parasites transfected with the centromeric plasmid presented an average reporter activity increase of 2.73-fold/day during eight days of drug selection (**Fig 3D, Supplementary Table 3**), greater than the rate of signal increase observed during selection of parasites transfected with the non-centromeric plasmid that also express *hdhfr* under control of the *pvhsp70* promoter (average increase of 1.34-fold/ day, **Fig 3B, Supplementary Table 3**). Over a period of two weeks’ drug pressure, transgenic parasites were observed to have a growth rate similar to untransfected parasites (3 – 5-fold/day) and the NanoLuc signal increased linearly with the number of parasites **(Supplementary Table 4).** In serial dilution experiments, as few as 100 transformants could be detected by NanoLuc signal above background control from uninfected RBCs **(Supplementary Table 4).**

### *P. knowlesi* transformed with a *P. vivax* centromere-containing reporter plasmid completes the parasite life cycle in rhesus macaques and *An. dirus* mosquitoes

In experiments to evaluate the ability of pvcen-pvhsp70-D-NanoLuc transformants to complete the full life cycle while expressing NanoLuc *in vivo* under no drug selection pressure, the parasites were inoculated into a splenectomized rhesus macaque (rhesus #1). Parasites were observed in Giemsa-stained thin blood films at day 3 after inoculation **(Fig 4A, Supplementary Table 5)**, after which parasitemia rapidly increased up to 2% at day 5. At midnight of day 5, when the rhesus parasitemia reached 1.23%, *An. dirus* B and *An. dirus* X mosquitoes (31) were fed on the infected monkey; the procedure was repeated the next night at a similar parasitemia of 1.20% **(Fig 4A; Feeds #1 and #2)**. Seven days after each feed, 16 – 20 mosquitoes were randomly selected from the individual batch for dissection and midgut isolation. As shown in **Fig 4B** and **Supplementary Table 6,** Feed #1 yielded a smaller number of infected mosquitoes (11 of 16 dissected) than Feed #2 (19 of 20 dissected). Oocyst counts per mosquito were also lower in Feed #1 than in Feed #2 (**Fig 4B**; 2.5 *vs*. 16.6 oocysts per mosquito on average, respectively). NanoLuc determinations from individual isolated midguts confirmed higher signals in proportion to the number of oocysts per midgut **(Fig 4C, Supplementary Table 6)** (R^2^ = 0.6903).

**Figure 4.**
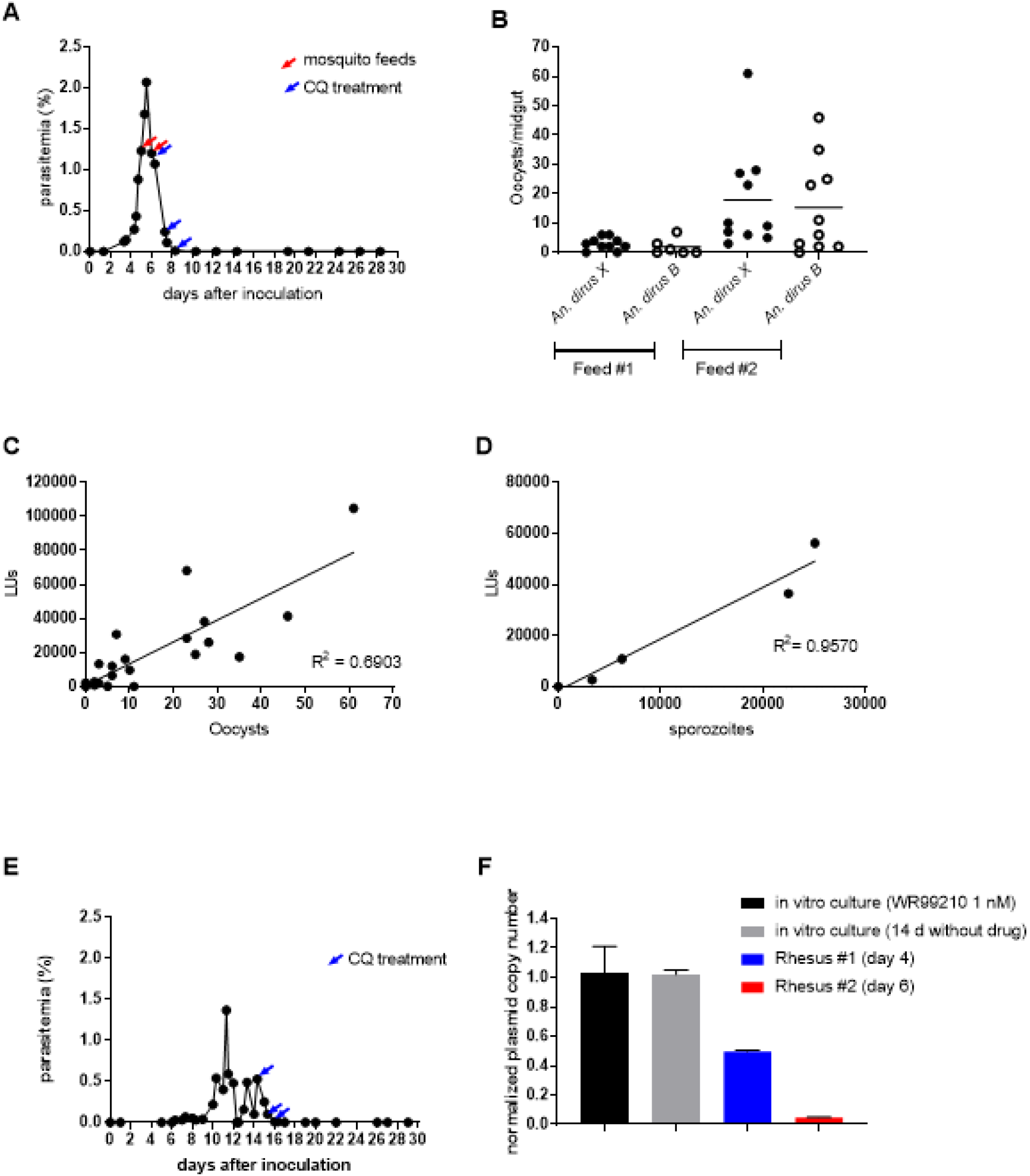
Bioluminescent *P. knowlesi* transformants complete the parasite life cycle *in vivo*. (A) Development of blood stage parasitemia in a splenectomized non-naïve rhesus macaque (rhesus #1) infected with pvcen-pvhsp70-D-NanoLuc-transformed *P. knowlesi*. The red arrows indicate when mosquito feedings were performed; blue arrows indicate administration of chloroquine (50 mg/kg, oral). (B) Counts of oocysts in the mosquito midgut 7 days after blood feeding. (C) NanoLuc activities from midguts isolated from Feed #2 infected mosquitoes. (D) NanoLuc activities from sporozoites isolated from mosquitoes 10 days after either Feed #1 or Feed #2 on rhesus macaque DCID. (E) Blood stage parasitemia developed in a splenectomized non-naïve rhesus macaque (rhesus #2) after bites of mosquitoes carrying infectious sporozoites. The blue arrows indicate chloroquine treatments (50 mg/kg, oral). (F) Copy numbers of the NanoLuc coding sequence relative to those of the single-copy *P. knowlesi aldolase* gene in blood-stage parasites from *in vitro* cultures and from blood samples of rhesus monkeys #1 and #2. Copy number results from each sample are presented relative to the copy number of pvcen-pvhsp70-D-NanoLuc-transformed *P. knowlesi* parasites cultivated under WR99210 selection pressure (black): cultivated parasites maintained 14 days *in vitro* without drug pressure (gray); parasites from rhesus #1 (blue); parasites from rhesus #2 (red). Error bars represent standard error of the mean; LUs, Luminescence units.

At day 10 after the feeds, eight mosquitoes from each batch were randomly selected and dissected for salivary glands and sporozoite detection. Total sporozoites from each group were counted, and, as expected from the oocyst data, higher sporozoite numbers were observed in mosquitoes from Feed #2 than Feed #1 (29,688 *vs*. 5,953 sporozoites per mosquito on average, respectively); no significant difference was found between the *An. dirus* B and *An. dirus* X strains **(Supplementary Table 7)**. NanoLuc activity was correspondingly higher from the greater number of sporozoites in Feed #2 mosquitoes **(Fig 4D; Supplementary Table 7)** (R^2^ = 0.9516).

The remaining sporozoite-infected mosquitoes were used to infect a second non-naïve macaque (rhesus #2). Six days after the mosquito bites and sporozoite injection, parasites were detected in Giemsa-stained thin blood films of peripheral blood **(Fig 4E; Supplementary Table 5).** Development of the infection was similar to previous reports of rhesus infections using sporozoites (via mosquito bites) (31,42). NanoLuc activity was readily detected in blood samples, even when taken at a low 0.03% parasitemia (data not shown).

### Centromeric plasmids are present at reduced number in transformed parasites that complete the full life cycle without drug selection

During the course of rhesus macaque infection #1, transmission to mosquitoes, and infection of rhesus macaque #2, the pvcen-pvhsp70-D-NanoLuc transformants were not subjected to drug pressure. To assess the plasmid maintenance over the life cycle in this absence of drug selection, including mitosis during the asexual parasite multiplication and meiosis after zygote formation, DNA was extracted from parasite samples and qPCR analysis was performed to quantify copy number of the NanoLuc cassette relative to the endogenous chromosome copy of the *aldolase* gene. The results indicated plasmid copy number loss during *in vivo* infections: parasites isolated from rhesus #1 at infection day 4 presented only half of the plasmid copy numbers of the parasites from *in vitro* cultures maintained for 14 days without drug pressure (**Fig 4F, Supplementary Table 8**). Parasites isolated from rhesus #2 at infection day 6, showed a 20-fold plasmid copy number loss compared to parasites from the unpressured *in vitro* culture. In marked contrast, parasites from the *in vitro* cultures showed no plasmid copy number loss relative to the *aldolase* gene after 14 days without drug pressure **(Supplementary Table 8)**.

### NanoLuc facilitates high-throughput assays of anti-malarial compounds and parasite invasion

To evaluate for potential use of NanoLuc-transformed *P. knowlesi* line in assays of antimalarial compounds, IC_50_ drug response measures assessed by NanoLuc signals were compared to the measures from standard SYBR Green assays. Results were obtained from transgenic and non-transgenic (“wild-type”) parasites exposed to chloroquine, WR99210, and G418 **(Table 1, Supplementary Table 9).** No significant differences were observed between the IC_50_ values from the two methods, and *P. knowlesi* transformants presented chloroquine and G418 IC_50_ values similar to untransfected parasites **(Table 1, Supplementary Table 9).** WR99210 IC_50_ values of the transformants were about 100-fold higher, consistent with *hdhfr* expression from the pvcen-pvhsp70-D-NanoLuc plasmid.

**Table 1.**
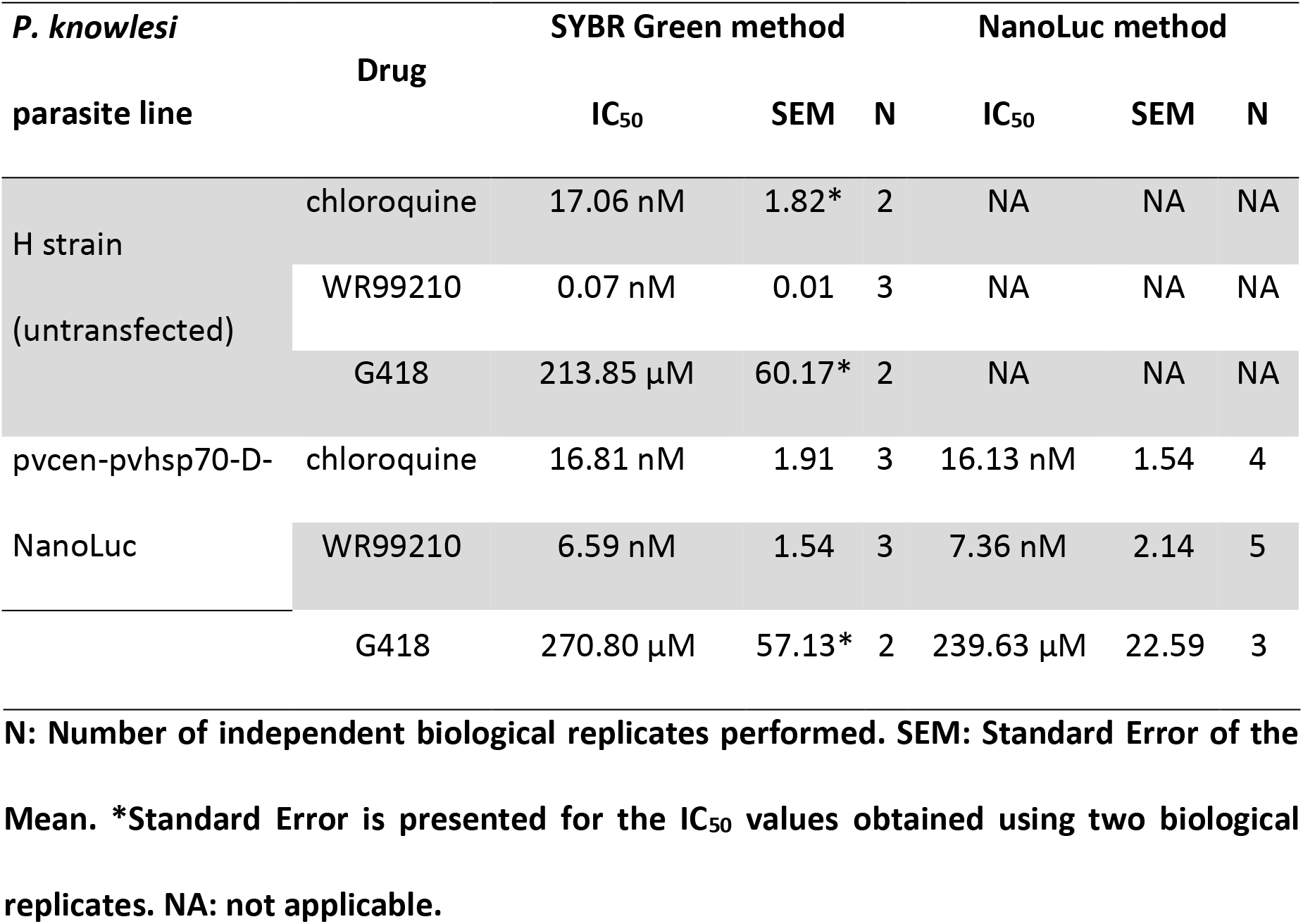
Antimalarial half-maximum inhibitory concentration (IC_50_) values obtained by SYBR Green and NanoLuc assays.

Invasion assays were also performed using NanoLuc-transformed parasites. Tightly synchronized *in vitro*-cultivated *P. knowlesi* schizonts were used to invade RBCs from different animal sources – rhesus macaque, human, *Aotus nancymaee, Saimiri boliviensis* and mouse. **Table 2** and **Supplementary Table 10** show that no significant difference in the results was observed between the Giemsa-stained thin blood film counts and the luminescence activity of the NanoLuc-transformed parasites, confirming that luminescence can be used to evaluate invasion of RBCs.

**Table 2.** *Plasmodium knowlesi* pvcen-pvhsp70-D-NanoLuc red blood cell invasion, determined by microscopy and NanoLuc activity.

|  | Microscopy |  |  | NanoLuc method |  |
| --- | --- | --- | --- | --- | --- |
| RBC source | RBC invasion | SEM | N | RBC invasion | N |
|  | relative to rhesus |  |  | relative to rhesus |  |
|  | RBCs |  |  | RBCs |  |
| Human | 0.74 | 0.11 | 3 | 0.84 | 1 |
| *Aotus nancymae* | 0.53 | 0.04 | 3 | 0.34 | 1 |
| *Saimiri boliviensis* | 0.34 | 0.01 | 3 | 0.40 | 1 |
| mouse | 0.03 | 0.02 | 3 | 0.06 | 1 |

**N: Number of independent biological replicates performed. SEM: Standard Error of the Mean.**

## Discussion

Previous evaluations of *P. vivax* regulatory sequences in *P. falciparum* showed that many were inactive (22), suggesting significant differences in the utilization of promoter elements by these species. The *P. vivax* 5’ UTRs studied in this work functioned in *P. knowlesi* even though some were inactive in *P. falciparum*. The 5’ UTRs of *pvcam* and *pvcrt* are two examples of flanking regions capable of driving the fLuc in *P. knowlesi* but not in *P. falciparum*. In contrast, the 5’ UTR of *pvhsp70* has activity in both parasites, driving at least 4-fold greater expression than the control *pfcam* 5’UTR. The longer length of the *pvhsp70* 5’UTR (1.4 Kb) may contribute for this stronger promoter activity. Highly conserved promoter elements in the *P. vivax* heat-shock gene evidently support similar levels of transcription in transformed *P. knowlesi* and *P. falciparum* parasites. In this regard, the presence of five copies of a GCATAT element in the 5’ UTR of *pvhsp70* may be relevant as this element has been identified as a feature of some deeply conserved genes in divergent branches of *Plasmodium* evolution (22). Absence of GCATAT from the 5’ UTRs of *pvcam* and *pvcrt* is consistent with less conserved promoters that are operative in both *P. knowlesi* and *P. vivax* but are not recognized in *P. falciparum*.

Episomal plasmids replicate by a rolling-circle mechanism leading to the formation of concatemers, resulting in variable copy numbers of the plasmid (43) and wide ranges of reporter expression. Furthermore, while episomal plasmids are useful tools for genetic studies; their maintenance requires drug pressure to prevent plasmid loss during cell division. To obviate these difficulties with copy numbers and generate transgenic lines that express reporter sequences through the complete parasite life cycle, plasmids containing centromeric sequences have been developed and used in *P. berghei, P. yoelii, P. falciparum*, and *P. cynomolgi* (34,40,43,44). Previous reports showed transgenic parasites containing centromeric plasmids were selected faster than parasites transfected with regular episomal constructs (34,40). Therefore, in the present study cultivated *P. knowlesi* parasites were transfected with the pvcen-pvhsp70-D-NanoLuc plasmid that contains *P. vivax* centromeric sequences and an *hdhfr* selection cassette. The resulting parasites expanded more than twice as fast as transformants containing non-centromeric plasmids, consistent with improved segregation and maintenance of the centromeric plasmid. Expression of the ultra-bright NanoLuc reporter sequence was confirmed throughout the *in vivo* life cycle, in blood-stage parasites, oocysts and sporozoites, and this expression was linearly related to the number of parasites in the assays. This linearity is an advantage of centromeric plasmids, that has been exploited successfully in mosquito stages, allowing its use in studies of mosquito-stage gene control and transmission blocking strategies (40). However, without drug selection, the episomal plasmid showed copy number loss over the life cycle, particularly *in vivo*. This finding highlights an advantage of transgenic lines that contain reporter sequences integrated into chromosomes of the parasite genome, achievable for example by recently developed zinc-finger gene modification methods (45,46) or by CRISPR/Cas9 technology (47,48). Whether the persistence of centromere-containing plasmids in *P. knowlesi* can be improved by use of homologous *pkcen* instead of *pvcen* elements remains to be investigated.

Rapid assay determinations and sensitivity allowing the use of low parasite numbers are among the advantages of NanoLuc-expressing parasites for high-throughput screening platforms. In the present work, NanoLuc expressing *P. knowlesi* were successfully used in drug response tests of blood-stage parasites *in vitro*, and the IC_50_ values obtained by luminescence assays were comparable to IC_50_ values obtained by the SYBR green method. Additionally to their use in drug response, centromeric plasmids such as pvcen-pvhsp70-D-NanoLuc may be useful in other experimental applications such as in vivo imaging. The *P. knowlesi* transformants can also be useful for validation of plasmid constructs prior to their use in transfection of *P. vivax*, which requires expensive non-human primate resources. The ultrabright and stable NanoLuc reporter has been previously used in *P. falciparum* (49) and *P. berghei* (29,50), producing luminescence up to 150-fold higher than firefly luciferase, and enabling detection of parasites at low densities and as single sporozoite (50). Recently developed methods of mosquito infection from *in vitro P. knowlesi* cultures (11) may thus provide useful strategies in high-sensitivity screens for transmission blocking vaccines and inhibitors of liver-stage parasites (29,49,50).

## Conclusions

*P. knowlesi* parasites can drive expression *P. vivax* promoters that are poorly recognized by *P. falciparum*, consistent with the closer evolutionary relationship of *P. vivax to P. knowlesi* than to *P. falciparum*. However, more conserved promoters like that of *pvhsp70* retain similar recognition by *P. vivax* and *P. falciparum*. Centromeric sequences from *P. vivax* can improve the efficiency of genetic transformation and may promote the stability of transfected plasmids across the complete *P. knowlesi* life cycle, through rhesus monkeys and *An. dirus* mosquitoes. NanoLuc-expressing, centromere-stabilized plasmids may be useful for high-throughput antimalarial screenings as well as biological imaging of *P. knowlesi* in the mosquito and vertebrate hosts.

## Declarations

### Ethics approval and consent to participate

All Non-human primate care and use in this study were performed in accordance with the NIH Animal Research Advisory Committee (NIH ARAC) Guidelines, under an Animal Study Proposal approved by the NIAID Animal Care and Use Committee (NIAIDLMIV 9E).

### Consent for publication

Not Applicable

### Availability of data and materials

All data generated or analyzed in this study are provided in the main text of this published article or as additional files.

### Authors’ contributions

RMB, JSA, JMS, JHA, OK and TEW designed the study. RMB, KT, JSA, TJG, RLC, MK, WAK, JPM, JKB, TE, LL, SOG performed experiments. RMB, JSA, TJG, JMS and TEW wrote the manuscript. All authors read and approved the final manuscript.

## Acknowledgements

We thank Patrick E. Duffy for providing the *P. knowlesi* H strain, rhesus blood samples for *in vitro* cultivation of parasites, and animal infections; Billeta Lewis, Milton J. Herrera, and Kelly E. Dicken, for monkey care; and Andre Laughinghouse and Kevin L. Lee, for providing and maintaining mosquitoes.

## Additional Files

**Additional file 1.**

Additional_file_1.xls

Supplementary tables

Supplementary Table 1. Oligonucleotide primers used in this work

Supplementary Table 2. Transient transfection of *Plasmodium* parasites

Supplementary Table 3. *Plasmodium knowlesi* growth and luminescence signal during selection using WR99210

Supplementary Table 4. NanoLuc signal of *in vitro* cultivated transgenic blood-stage parasites

Supplementary Table 5. Rhesus infections with transgenic luminescent *P. knowlesi*

Supplementary Table 6. Number of oocysts detected in individual mosquitoes dissected 7 days after the feeds and Luminescence signals obtained (only midguts isolated from mosquitoes from Feed #2)

Supplementary Table 7. Total sporozoites isolated from 8 mosquitoes from each species, infected with transgenic bioluminescent *P. knowlesi*, 10 days after the feeds and Luminescence signals

Supplementary Table 8. pvcen-pvhsp70-D-NanoLuc plasmid copy number variation in blood-stage parasites

Supplementary Table 9. Half-maximum inhibitory concentration (IC_50_) measures of drug response

Supplementary Table 10. *Plasmodium knowlesi* red blood cell invasion assays

